# Systemic diazepam alters local hippocampal CA1 circuits and differentially affects entorhinal and CA3 inputs

**DOI:** 10.64898/2026.08.18.745488

**Authors:** Jen G. Peterson, Matthew T. Erickson, Ayaka Sheehan, Chelsey C. Damphousse, A. David Redish

## Abstract

The GABA_A_ positive allosteric modulator diazepam is taken systemically by millions of people daily. GABA_A_ signaling is essential for hippocampal circuit function, but the effects of systemic diazepam on hippocampal information processing during behavior has not been studied. To answer this question, large neural ensembles were recorded from rats running a linear track under systemic diazepam administration. A cross-correlation of spiking activity revealed significantly increased inhibition from interneurons, aligned with the timescale of GABA_A_, suggesting a direct effect on local circuits. Local field potentials (LFP) showed an increase in theta and lo-gamma (30-50 Hz) power but a decrease in hi-gamma (80-120 Hz) power. We also found decreased amplitude and rate of sharp wave ripple (SWR) events and a reduction of firing rate and proportion of cells recruited to the SWRs. An autocorrelation of single-cell spike trains revealed a decrease and shift from shorter to longer timescales, aligning differently with theta frequencies. Phase coupling measurements showed decreased cellular coupling to theta and increased coupling to lo-gamma and hi-gamma. Finally, entropy of decoding along the track was increased, suggesting less precise spatial representations under diazepam. These changes suggest mechanisms that would likely disrupt hippocampal memory storage and consolidation processes under systemic diazepam.

## Introduction

Diazepam is a positive allosteric modulator (PAM) which increases the efficacy of GABA_A_ receptors, particularly through the α1-3, and α5 subunits ^1^. Due to this, diazepam is expected to have profound effects on neural activity and circuits depending on GABA_A_ functioning. Information processing in the hippocampus is highly dependent on inhibition through the GABA_A_ receptor, both internally in hippocampus ^2–5^ and through inputs from other structures, including medial septum ^6–9^ and prefrontal cortex ^10^, particularly through the nucleus reunions ^11^. Previous research has explored the effects of diazepam locally on the hippocampus in vivo ^12^ and in vitro ^13,14^. However, clinically, diazepam is taken systemically, not locally injected into the hippocampus. Therefore, it is important to investigate how systemic delivery of diazepam affects hippocampal functions and information processing within the neural circuits of CA1.

Diazepam is known to have memory-related side effects in practice, but the underlying causes of those memory-related side-effects are still poorly understood ^15–17^. Previous work looking at diazepam in behavior in rodents has found that diazepam impairs the formation of new memories (anterograde amnesia), memory consolidation (recent retrograde amnesia), but not the retrieval of well-consolidated memories ^18–21^. Recent studies have found that systemic diazepam affects hippocampal processes, known to be involved in memory storage, recall, and consolidation. In particular, diazepam disrupts *sharp wave ripple complexes (SWR)*, including reducing the power, amplitude, number and duration of SWR events ^14^. It also reduces reticular-elicited hippocampal theta frequency ^22^.

Both hippocampus and the inputs to hippocampus that are critical to its function (such as medial septum and medial prefrontal cortex through nucleus reunions) have GABA_A_ receptors likely to be differentially affected by diazepam. The GABA_A_ receptor subunits that diazepam acts upon are distributed heterogeneously in the layers of the hippocampus ^14^ and throughout these circuits. The subunits are distributed in different layers, so diazepam acts differentially on GABA_A_ receptors throughout the hippocampus ^23^. While this may explain its varied effects on the hippocampus, the changes driven by diazepam have not been studied, either in terms of its effects on hippocampal layers or its effects on place cell function.

The hippocampus typically demonstrates two processing states: *theta*, characterized by strong power in the 6-10 Hz theta band, and *LIA (Large-amplitude Irregular Activity)*, characterized by broader spectral signals, punctuated by transient 140-200 Hz SWRs ^6,24–27^. These two processing states are used to create and update a “cognitive map” which allows for spatial representation of the world ^6,24^. Pyramidal neurons within the CA1 reveal themselves as place cells behaviorally — each cell fires in a limited portion of an environment (the place field), and it is possible to decode representations of spatial locations from the distribution of active cells ^28,29^. During each theta cycle, the decoded location first encodes the location of the animal and then sweeps forward down potential paths ^30,31^ running to the next goal in a sequence ^32–34^. During LIA and SWRs, the cells fire sequences consistent with the structure of the environment ^35–38^. Hippocampal SWR and replay states are critical components of memory consolidation ^39–43^.

In this study, we report effects of systemically-delivered diazepam on hippocampal function from two experiments. First, we report changes in single cell and local field potential (LFP) patterns from multi-shank silicon probes recording at the pyramidal layer from rats running a simple linear track. Second, we reanalyze layer-specific LFP data from a previous experiment recording from linear silicon probes from rats running on an approach-avoidance motivational conflict task. We find profound disruptions in hippocampal function in both theta and LIA states likely to have strong implications for spatial navigation, memory, and consolidation processes.

## Results

Silicon probes implanted in dorsal hippocampal CA1 were used to record neural activity while rats ran on a linear track for food reward. Diazepam (1-3 mg/kg) or vehicle control was injected systemically (intraperitoneal) 10 minutes prior to running. Rats had 30 minutes to run for food on the linear track and then rested in a separate enclosure for 20 minutes of post-task recording. Rats underwent a dose sequence alternating control and diazepam doses (1-3 mg/kg). Prior to the dose sequence, rats typically had 1-2 weeks of training on the linear track, where they learned to run back and forth between two feeders providing 45mg food reward each (Fig. 1A).

**Figure 1.**
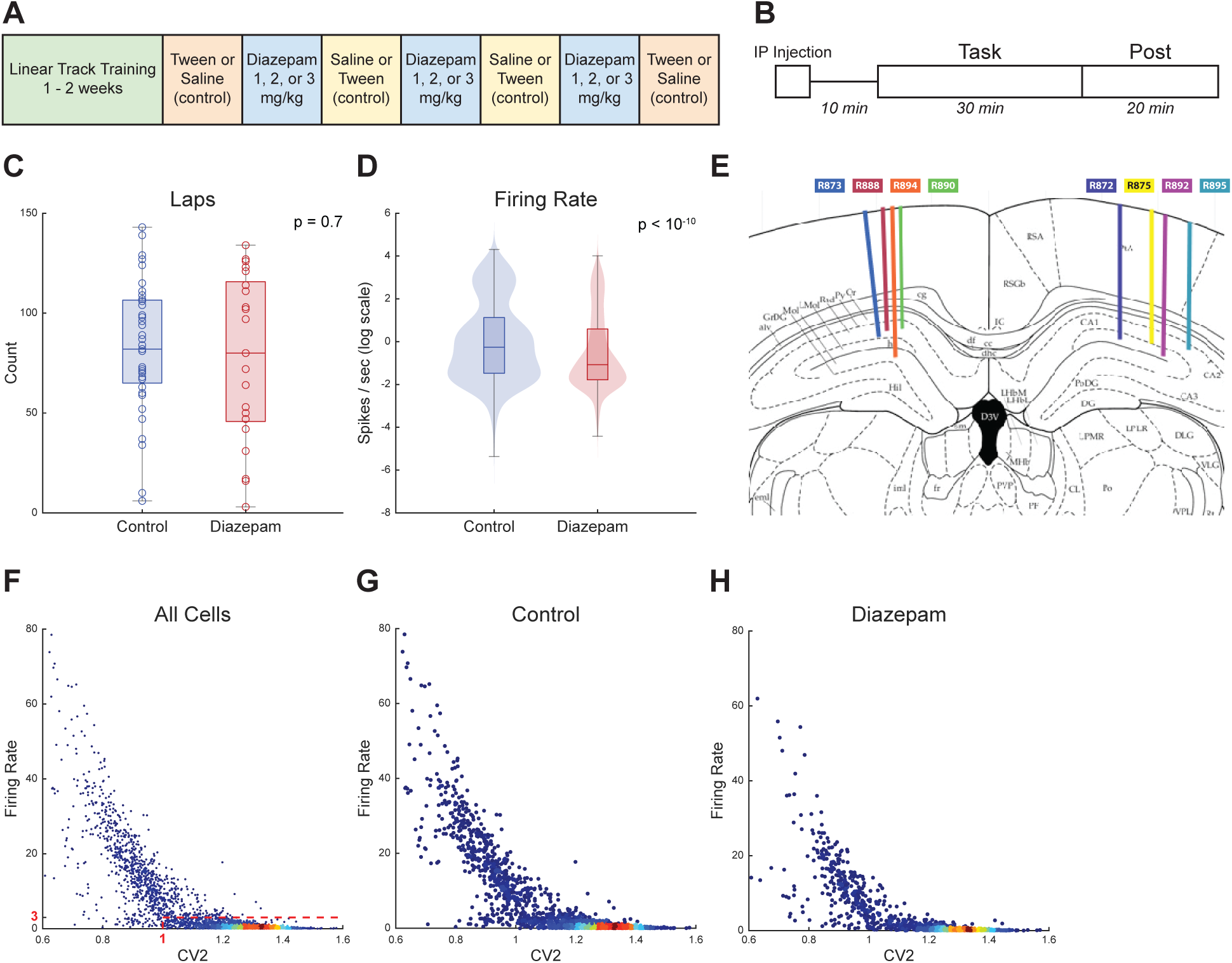
(A) Experimental sequence. Controls were alternated between saline or tween, either tween, saline, saline, tween or saline, tween, tween, saline. Each rat received 1, 2, or 3 mg/kg in random order. (B) Timeline of an individual session. Rats received IP injections 10 minutes before the task which lasted for 30 minutes, followed by a post-task recording in a covered pot. (C) Number of laps rats ran on average for control and diazepam conditions. (D) Firing rate distribution of cells for control and diazepam conditions. (E) Histologically identified probe placements. Each color represents the placement of the probe in an individual rat. (F-H) Distribution of coefficient of variance of interspike intervals (CV2) and firing rate over individual cells used to differentiate pyramidal cells from interneurons. (F) Combined distribution over all sessions. The dashed red box represents the cutoffs for interneuron or pyramidal cells as defined by firing rate and CV2. (G) Distribution over control sessions. (H) Distribution over diazepam sessions.

Rats ran a similar number of laps on control days as diazepam days. To increase sample size and statistical sensitivity, controls (no drug, saline, tween) and diazepam doses (1-3 mg/kg) are each combined for analysis. Comparing combined control and diazepam dose data found no significant difference between the number of laps rats ran (Mann-Whitney non-parametric test, p = 0.71). However, when analyzed separately, higher doses of diazepam (2 mg/kg and 3 mg/kg) showed a significant decrease in laps (ANOVA, p = 0.0003, d.f. = 5, F = 5.51), see supplemental figure (Fig. S1A). Laps were normalized by rat to account for differences between individual animals (Fig. 1B). The firing rate of all recorded cells under diazepam was significantly lower than controls (Mann-Whitney non-parametric test, p < 10^-10^). Higher doses of diazepam also resulted in lower overall firing rates, showing a dose-effect (Supplemental Figure S1B, ANOVA, p < 10^-16^, d.f. = 5, F = 15.62). (Fig. 1C).

For later statistical analyses, interneurons and pyramidal cells were separated by thresholds of firing rate and the coefficient of variation of the interspike intervals (CV2) ^2,3^. As can be seen in Fig. 1D, the distribution of cells separated into two clusters on these two dimensions, consistent with extensive historical recordings in hippocampus ^24,48,49^. Following previous determinations, cells with CV2 > 1 and firing rate < 3 Hz were identified as putative pyramidal neurons, while cells with CV2 < 1 or firing rate> 3 Hz were identified as putative interneurons. Because the distributions were similar under diazepam and control, we used these same thresholds to identify putative cell type under diazepam and control (Fig. 1D).

GABA_A_ receptors are found on both interneurons and pyramidal cells in the hippocampus ^23^. Diazepam enhances inhibition by increasing the frequency of channel opening ^50^, acting as a GABA_A_ positive allosteric modulator, and increasing the efficacy of GABA_A_ functionality. Given that the timescale of GABA_A_ receptors is around 10 ms ^51^we expect that diazepam should result in increased hyperpolarization and less cell firing 10 ms after inhibitory cell spiking.

To directly measure the synaptic effects of one cell on another, we cross-correlated spiking activity for simultaneously recorded pairs of cells. Using the identified cell types of pyramidal and inhibitory neurons (Fig. 1D), cell pairs were separated into four combinations: pyramidal to pyramidal, pyramidal to inhibitory, inhibitory to pyramidal, and inhibitory to inhibitory. Cross-correlations between inhibitory neuron pairs showed a decrease in probability of spiking in the postsynaptic cell approximately 10 ms relative to the reference spike (output) of the presynaptic inhibitory neuron, consistent with an increase in GABA_A_ efficacy (Fig. 2D). We found a smaller but similar effect in inhibitory to pyramidal pairs (Fig. 2C). Importantly, no significant effects were found between pyramidal pairs or between pyramidal to interneuron pairs, confirming the fidelity of the GABA_A_ effects of diazepam. Similar effects were seen in the interneuron pairs (Fig. 2H). However, during the post behavior recording, pyramidal to pyramidal inputs also showed a decrease in the probability of spiking around 5 ms (Fig. 2E). This may be due to the reduction in SWR events and spiking during SWRs as discussed below, and may be due to a reduction in the increase of cell-coupling often seen after experience ^52–54^.

**Figure 2.**
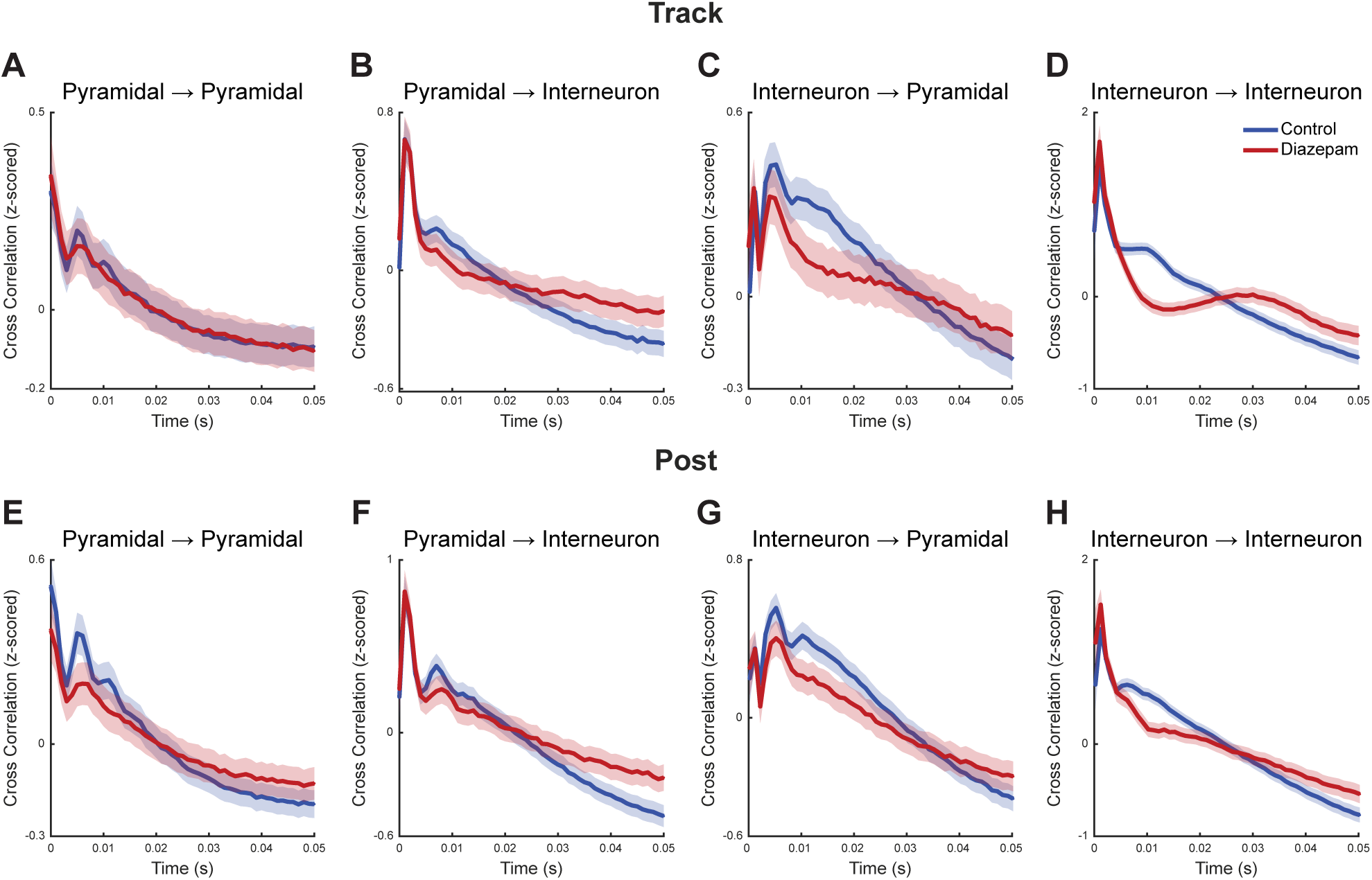
Z-scored cross correlation between cell pairs reveals synaptic effects. In each panel, the blue line shows control sessions, and the red line diazepam sessions. (A-D) effects from the 30 minute on-track recordings: (A) pyramidal → pyramidal, (B) pyramidal → interneuron, (C) interneuron → pyramidal, (D) interneuron → interneuron. (E-H) Equivalent plots from the 20 minute *post-run* recordings.

LFPs were also recorded and found to change under pharmacological manipulation. Consistent with the very large hippocampal literature, we found that the task portion of the recording was dominated by the theta state and the post recording by LIA (Fig. 3) ^24,27^.

**Figure 3.**
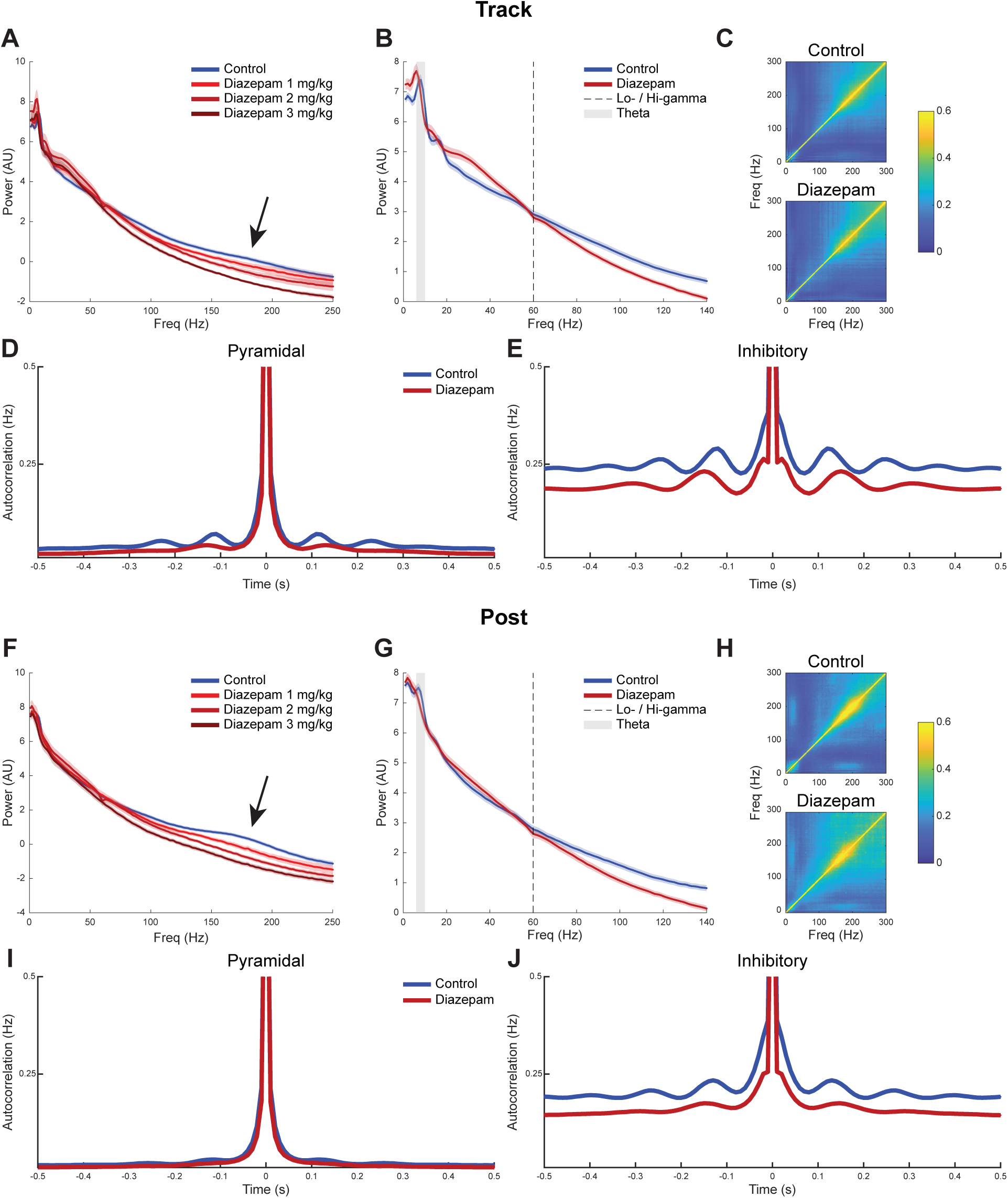
Spectral analyses. Power spectral densities on the track and in the post-run recording. (A) PSD shown out to 250 Hz, including differentiated doses to show the dose-response curve in the SWR band from data recorded on the track. (B) PSD shown out to 140 Hz to show changes in theta (6-10 Hz), lo-gamma (30-50 Hz), and hi-gamma (80-120 Hz) from data recorded on the track. The black arrow indicates the sharp wave ripple band. The dashed line shows the cut-off between high and low gamma (C) Cross-frequency plots revealing SWR events on the track. (D-F) parallel analyses performed from data from the post-run recording. Spike autocorrelation analyses. (F) Pyramidal neurons on the track. (G) Interneurons on the track (C) Pyramidal neurons during post-run rest. (D) Inhibitory neurons during post-run rest.

Power spectral densities (PSD) are most sensitive to constant or rhythmic frequencies like theta (6-10 Hz) and the various gamma signals typically seen in CA1 (lo-gamma, driven by CA3 inputs 30-50 Hz and hi-range gamma, driven by entorhinal inputs 80-120 Hz) ^55–57^. In contrast, frequency-autospectrum cross-coherence plots examine the correlation between frequencies ^45^, and are more sensitive to momentary events such as SWRs (Fig. 3C). Coherence correlation plots are less sensitive to constant frequencies, thus theta and gamma do not appear as strongly. A PSD analysis revealed a frequency shift to lower frequencies in the theta range under diazepam both on the track (Fig. 3A) and in the post-run recording (Fig. 3B), suggesting a slowing of theta oscillations under diazepam. Additionally, we found an increase in power of lo-gamma (30-50 Hz) and a decrease in power of hi-gamma (80-120 Hz) under diazepam both on the track and in the post-run recording.

Under all doses of diazepam, LFP showed a power reduction in the 140-200 Hz SWR frequency range (Fig. 3A, D, indicated by black arrow). The effect was present both on the track, and in the post-run recording, but appeared stronger in the post, likely due to the rats being more commonly in a LIA state in the post-run recording where SWRs occur more frequently. However, we did see a reduction in SWR power on the task as well, likely attributable to the diminishment in SWRs during pauses at each feeder. A dose relationship was observed — larger doses of diazepam produced more reduction in the SWR power (Fig. 3A, F).

To confirm the diminishment of SWR events, not just power in the SWR frequency range, we measured cross-frequency auto-correlation plots ^45^. Correlation plots found a large band of correlated frequency in the SWR range. Compared to control, the SWR band was reduced under diazepam both on the track and in the post-run recording (Fig. 3B,G).

LFPs arise from synaptic events, which include signals from inputs, local circuits, and volume conduction ^26^. In order to identify whether these oscillatory changes were also appearing in the cell firing, we directly examined spike train autocorrelations. Cells were separated by pyramidal or interneuron per above. Under diazepam, oscillations revealed in the autocorrelation decreased and slowed in both the pyramidal cells and the interneurons (Fig. 3D,E,I,J). Theta oscillations were more visible on the track (Fig. 3D) than in the post-run recording (Fig. 3I), consistent with differences in expectations due to proportions of theta and LIA under the two conditions. On the track, the reduction in frequency was observed in both pyramidal (Fig. 3D) and inhibitory neurons (Fig. 3E), but was strongest in the inhibitory population. Additionally, a reduction in firing rate was observed in the inhibitory neurons as diazepam shifted the overall autocorrelation lower (Fig. 3J). No strong changes were observed in the pyramidal neurons during post behavior rest (Fig. 3I). The lack of effect may be due to the expected reduction in theta oscillations during the LIA state. However, the inhibitory neurons did reveal a similar effect to the track, with slowed frequency and lower overall autocorrelation.

Given the effects on power in the SWR frequency range and the effects on cell autocorrelation firing (Fig. 3), we examined the rate, amplitude, firing rate, and proportion of cells recruited to SWRs. We found that the rate (Fig. 4A,E) and amplitude (Fig. 4B,F) of SWRs, firing rate of cells during SWRs (Fig. 4C,G), and the proportion of cells recruited in each SWR (Fig. 4D,H) were all significantly reduced under diazepam in both the track (Fig. 4 A-D) and post (Fig. 4 E-H), indicating SWRs were severely impacted in both waking and resting states.The rate and amplitude of detected sharp waves was decreased under diazepam on the track (Mann-Whitney, rate p = 0.01, amplitude p < 10^-6^) and in the post-run recording (Mann-Whitney, rate p < 10^-10^, amplitude p < 10^-10^). The average number of spikes during SWR events fired by each cell decreased under diazepam on the track and in the post-run behavior rest (Mann-Whitney, task and post, p < 10^-10^). The proportion of cells recruited within each SWRs decreased under diazepam on the track and in the post-run behavior rest (Mann-Whitney, task and post, p < 10^-10)^.

**Figure 4.**
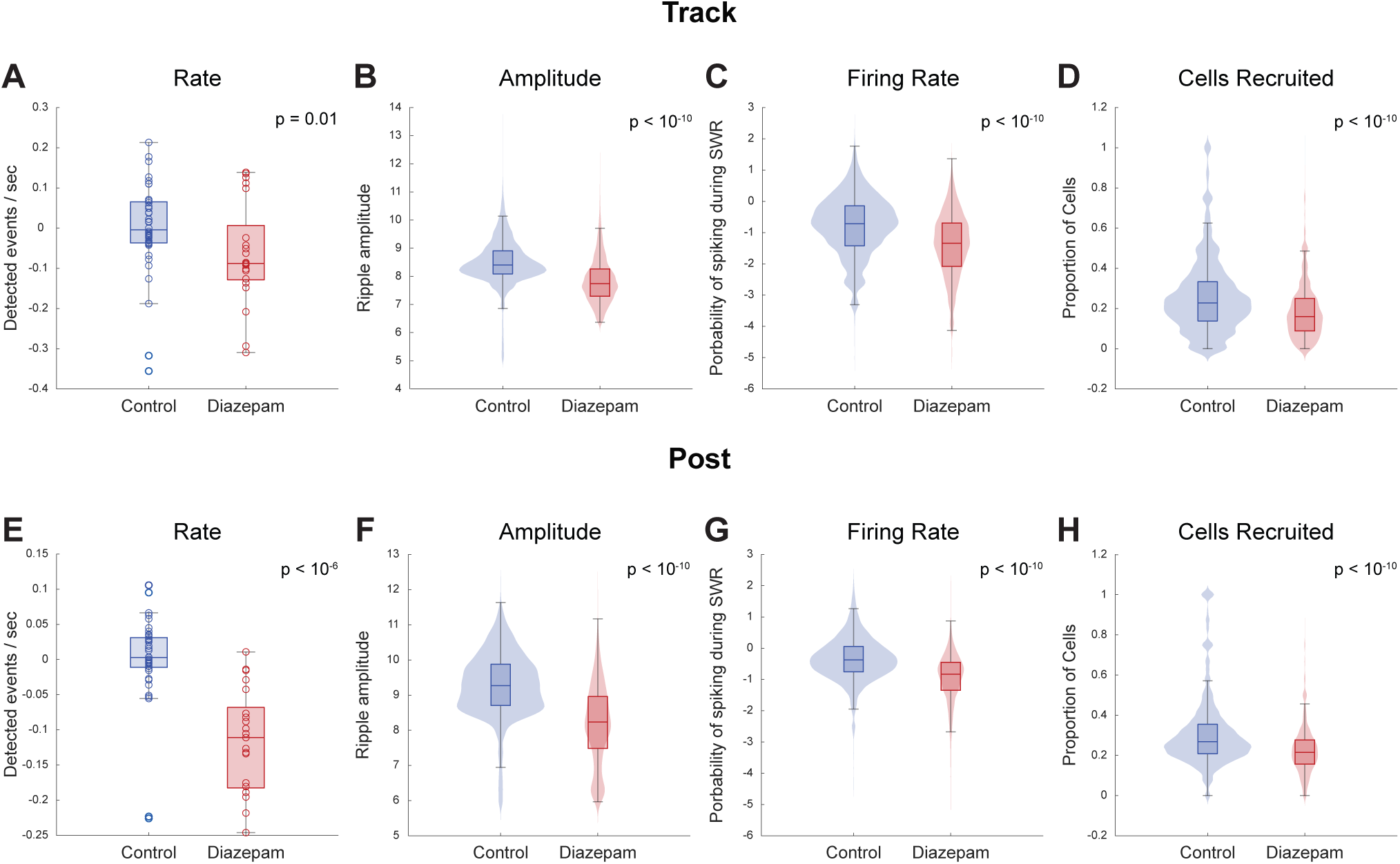
Sharp wave ripple (SWR) analyses. (A,E) SWR rate. (B,F) SWR amplitude. (C,G) Firing rate during SWRs. (D,H) Proportion of cells firing within a given SWR. (A-D) On the track; (E-H) During post-run recording.

CA1 receives inputs from other brain regions into different layers — medial entorhinal (MEC) inputs synapse onto the stratum lacunosum-moleculare and CA3 inputs onto the stratum radiatum. Observing effects throughout the layers may help us understand how systemic diazepam is changing these inputs.The use of longer probes in a previous cohort (originally reported in ^44^) allowed for recording of LFP but not cells from the multiple layers of CA1. These rats received either 2 mg/kg diazepam or Tween-20 vehicle control while running on an approach-avoidance motivational conflict task. We performed novel analyses of these data to look at the differences in layers from this previous data set. Figure 5B shows probe placement for each rat. All probes except one (indicated by red x, excluded from analysis) crossed the pyramidal layer of CA1 (Fig. 5B). Thus n=5 rats were re-analyzed here. Cross-frequency auto-coherence plots were used to visually select channels with the strongest SWR band. These channels were identified as the stratum pyramidale depth as reference. PSDs and coherence plots were generated and then averaged by depth in relation to stratum pyramidale.

**Figure 5.**
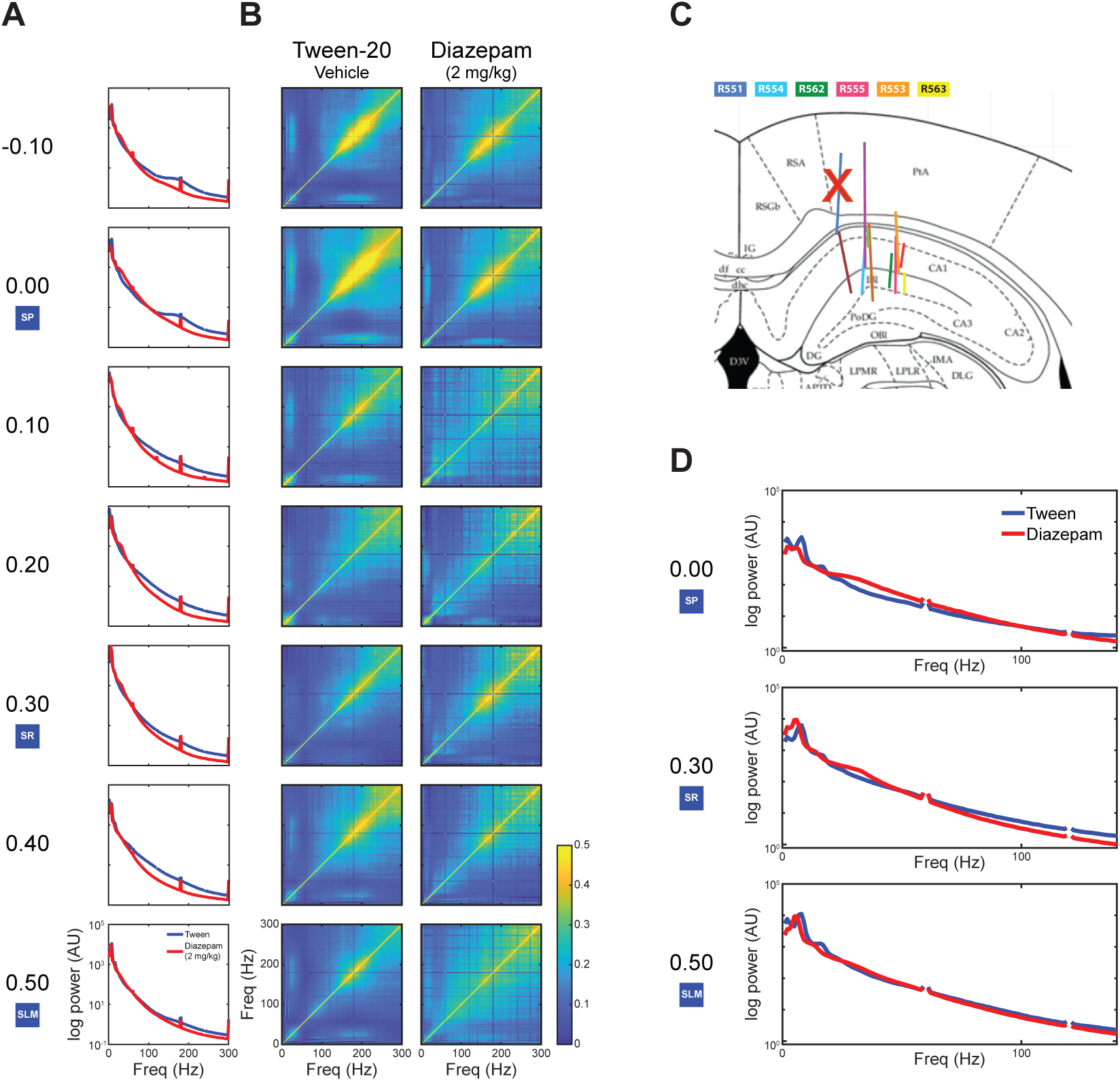
Novel analyses of the older data set from ^44^. (A) Power spectral densities and (B) cross-correlation plots by distance from the pyramidal layer. (C) Placement of probes in the second cohort based on histology. Red x represents the rat that was excluded from the dataset due to probe placement. (D) Zoomed in power spectral densities for each identified layer showing changes in theta (6-10 Hz), lo-gamma (30-50 Hz), hi-gamma (80-120 Hz). SP = striatum pyramidale (pyramidal layer); SR = stratum radiatum (CA3 inputs); SLM = stratum lacunosum-moleculare (entorhinal inputs). We have nan’d out 60 Hz due to the notch filter included in this data ^44^.

At the stratum pyramidale (0.00 mm), cross-correlation analyses revealed a reduction in the correlation of upper frequencies in the SWR band under diazepam (2 mg/kg), (Fig. 5A). PSDs also revealed a reduction in the SWR and theta power (Fig. 5A). Additionally, PSDs revealed an increase in lo-gamma power under diazepam (Fig. 5A, left panel). All of these observations are consistent with the primary cohort reported above (see Fig. 3).

In the stratum radiatum (+0.30 mm), we observed little correlation in the SWR band under either the control or diazepam. The PSDs revealed an increase in lo-gamma power and a decrease in hi-gamma power. Consistent with ripples being predominantly generated at the stratum pyramidale ^27^, we did not see an increase in power at the SWR band in the control as was observed in the stratum pyramidale. In the stratum lacunosum-moleculare (+0.50 mm), small SWR bands returned in the correlation plots under control conditions. Under diazepam, these bands were reduced throughout the whole SWR frequency range. In the PSDs, there was a decrease in the power of hi-gamma at the stratum lacunosum-moleculare.These data are consistent with a decrease in MEC connectivity, but a potential increase in CA3 connectivity under diazepam

Both pyramidal cells and interneurons fire at specific phases of the low-frequence LFP oscillations in the hippocampus, including theta, lo-gamma, and hi-gamma, and thus they show strong phase-coupling to these rhythms ^55–59^. To investigate how spiking changed relative to those LFP oscillations, we looked at phase coupling, using PPC (see Methods). PPC measures the average angular phase difference between all pairs of spikes and compares their z-scores relative to shuffled samples. Importantly, PPC is robust to both large and small samples ^46,47^. Because interneurons and pyramidal cells couple to LFP signals differently, phase coupling was calculated for theta, lo-gamma, and hi-gamma separately for interneurons and pyramidal cells.

Under diazepam, both interneurons and pyramidal cells were less coupled to theta (track: pyr (Fig. 6A) p<10^-7^, int (Fig. 6D) p<10^-4^, post: (Fig. 6G) pyr p<10^-31^, int (Fig. 6J) p<10^-14^). Fascinatingly, coupling to lo-gamma was increased for both cell types. (track: pyr (Fig. 6B) p<10^-22^, int (Fig. 6E) p<10^-40^, post: pyr (Fig. 6H) p<10^-3^, int (Fig. 6K) p<10^-16^). Much less of a change in coupling was observed for hi-gamma, although there was a statistically significant increase in coupling under diazepam for both cell types both on the track and in the post-run recording (track: pyr (Fig. 6C) p=0.03, int (Fig. 6F) p=0.0024, post: pyr (Fig. 6I) p<10^-6^, int (Fig. 6L) p=0.032). Under diazepam, cells were less coupled to theta and more coupled to lo-gamma, with much smaller changes relative to hi-gamma. CA3 inputs are associated with lo-gamma LFP frequencies, while MEC inputs are associated with hi-gamma LFP frequencies ^57^, suggesting one effect of diazepam is increased sensitivity to CA3 inputs, but less of a change to MEC inputs.

**Figure 6.**
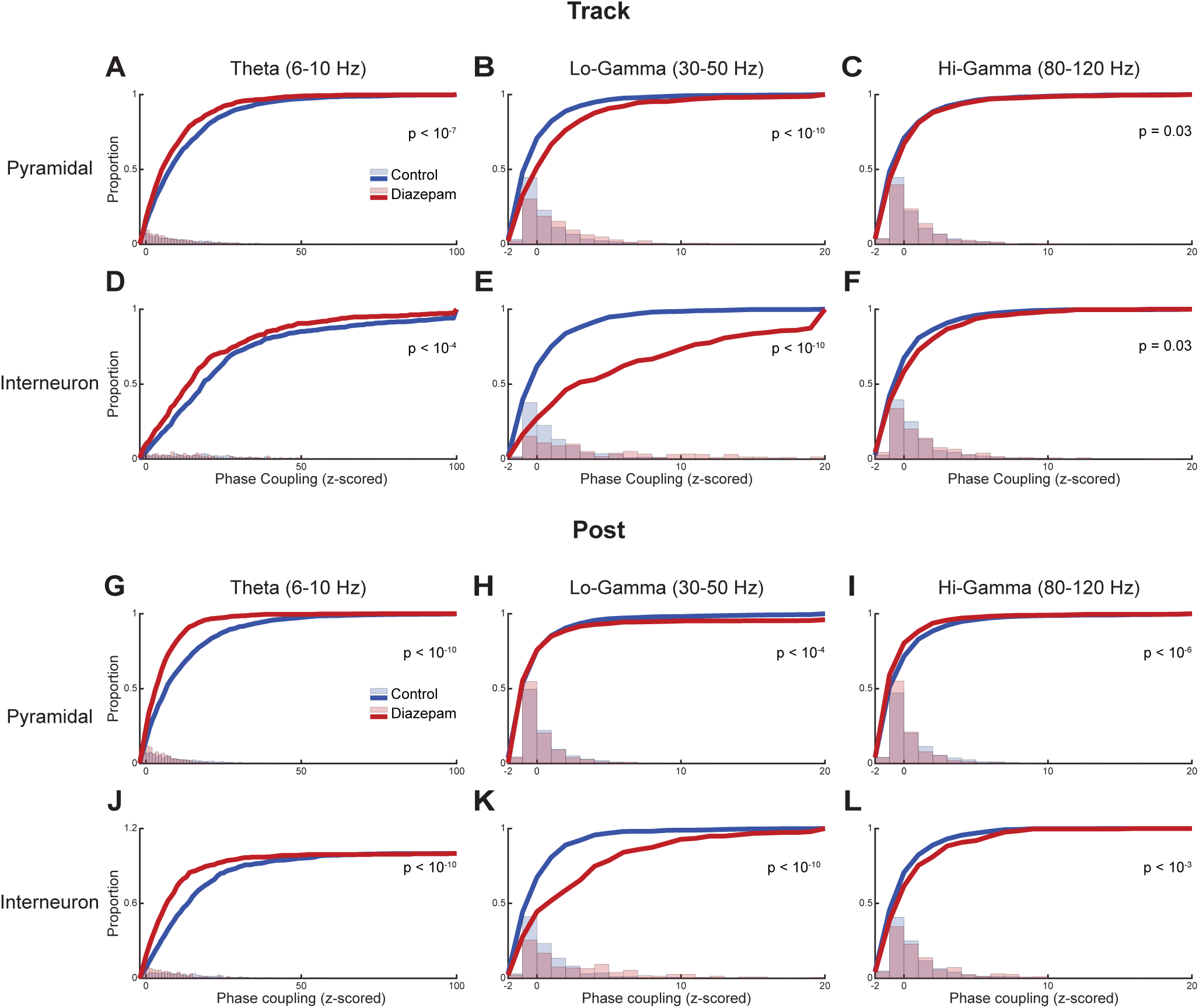
Phase coupling between cells and local field potential oscillations. Each panel shows the distribution of z-scored PPC values ^46,47^ in the bar plots and the cumulative sum (cumsum) plots in the solid lines. Bars and cumsum plots show the same data. *On the track*, pyramidal cells showed reduced coupling to theta (A), increased coupling to lo-gamma (B), and barely increased coupling to hi-gamma (C). Similar changes were seen in the interneurons (D, E, F). In the *post-run* recording, pyramidal cells showed decreased theta coupling (G), decreased lo-gamma coupling (H), and decreased hi-gamma coupling (I). Interneurons showed decreased theta coupling (J), increased lo-gamma coupling (K), and increased hi-gamma coupling (L).

These changes are likely to drive representational changes. In order to examine representational issues, Bayesian decoding was used to estimate location of the rat on the track based on cell firing using a one-step (history-less) decoding ^28,29^. Figure 7 shows two example sessions (one saline, Fig. 7A, and one diazepam [2 mg/kg] Fig. 7B). The red line displays the rat’s actual location and the cyan distribution shows the Bayesian posterior decoding of the position. Visually, the diazepam day has a “messier” decoding. To quantitatively measure the decreased reliability of the decoding, we measured the entropy of the posterior distribution for each decoded time bin. Entropy measures precision independent of accuracy. Entropy measured across all sessions from all rats showed a significant difference between controls and diazepam. The decoding entropy was significantly higher under diazepam both on the track and in the post-run recording (track, p=0.04, post, p=0.05), indicative of more representational noise under diazepam.

**Figure 7.**
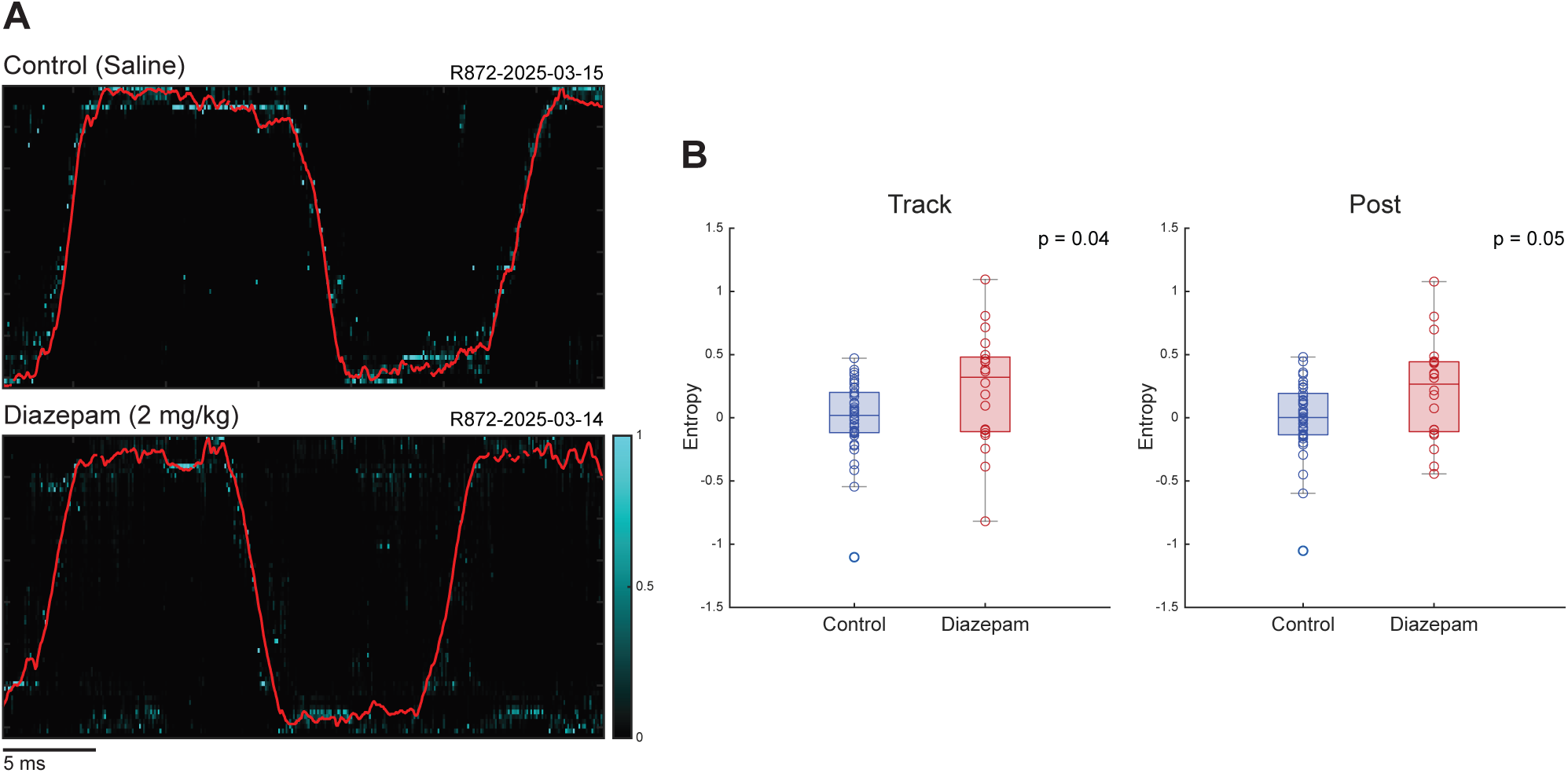
(A) Example decoding of location on the track for a control and a diazepam (2 mg/kg) day in the same animal. The position of the animal is shown in red, and decoding is shown in cyan, with brighter cyan indicating more decoding posterior. (B) Average entropy of all decoding samples on the track and during the post-run recording.

## Discussion

Systemic diazepam is given to millions of people daily^60^, and while there have been reports of memory deficits ^61–63^, how systemic diazepam changes memory function remains unknown. We tested a systemic dose of diazepam on hippocampal function and saw profound changes in hippocampal information processing, both in theta processing states andSWR events. In particular, even low doses of diazepam had devastating effects on SWR events, both during rest periods on the track (typically seen at feeder sites), and during post-run rest.

Theta rhythms are generated in the hippocampus via activity coordinated between the medial septum, local circuits, and entorhinal cortex, importantly, including GABAergic projections ^6^. Diazepam can be expected to impact these GABA projections resulting in the changes observed in oscillatory power, most specific in the frequency range of theta. Diazepam likely changes the efficacy of inhibitory neurons firing which could drive the frequency shift observed in theta under diazepam. Not only may local circuits be affected, but the timing and strength of the inputs could be affected, as systemic diazepam will include actions on other parts of the brain. Importantly, this means that if one wants to understand the effects of systemic diazepam, it must be studied through systemic distribution, not through local infusion. Because humans receive diazepam systemically, we chose to examine the effects of systemic provision.

A loss of the theta temporal structure in the hippocampus may alter sequence firings and thus corrupt memory consolidation of said sequences as mentioned above. Consistent with these hypotheses and previous observations that diazepam can affect hippocampal oscillations and cell coordination ^64,65^, we found changes both within the oscillations themselves (Fig. 3), in cell autocorrelation processes (Fig. 2), and in the coupling of cell firing to oscillations (Fig. 6).

Slow gamma oscillations arise through interactions with CA3 projections and fast gamma arises through interactions from medial entorhinal cortex ^57,59^, driving a *you-are-here, sweep-forward* representational sequence during behavior ^30,31,66^. The differential effects of systemic diazepam that we observed affecting slow gamma more than fast gamma suggest an impact on the *you-are-here* component driven by CA3 inputs more than the *sweep-forward* component. Fascinatingly, we found a decrease of cell coupling to theta oscillations, but an increase in cell coupling to slow gamma oscillations, suggesting an increased impact of CA3 inputs on CA1 under systemic diazepam (Fig. 5). The interplay between these two gammas and theta and how diazepam modulates this balance and the cell coupling to these frequencies may be responsible for the increased entropy we observed in our decoding measurements (Fig. 7).

SWRs are a key oscillatory component observed in the dorsal hippocampus that occur after rewards and during other non-attentive pauses while animals are on the track and are particularly prominent during post-task rest ^27,67^. They are modulated by a balance of excitatory and inhibitory circuits in CA1, and depend on phasic inhibition, primarily modulated by GABA_A_-mediated inhibitory post-synaptic potentials (IPSPs) ^27,68,69^. Thus it is not surprising that diazepam administration disrupted SWRs due to its profound effect on GABA_A_-mediated IPSPs. A large effect we observed was disruption of SWRs, including a loss of high-frequency power of the SWR LFP band, consistent with previous in vitro work ^12,14^. This effect is also consistent with previous in vivo studies manipulating GABAergic transmission more directly ^70,71^. Additionally, we found a reduction in rate, amplitude, firing rate during events, and proportion of cells recruited under diazepam.

SWRs have been hypothesized to play roles in memory consolidation, memory retrieval, and action planning ^24,27,35,72–75^. The largest effects are typically seen in post-task rest, where SWRs occur more frequently and support offline memory processes ^41,76–78^. The disruption of SWRs that diazepam causes likely impair these sequences playing during rest. In previous studies, disruption of post-task SWRs via electrical stimulation caused significant impairment on tasks requiring spatial memory ^41–43,79^. With its disruption, we can expect diazepam to have a profound effect on the memory processes associated with SWRs, particularly episodic memory consolidation, but these have not been extensively studied.

A possible explanation for the differential effects of diazepam, particularly on the differences between CA3 and entorhinal impacts on CA1 cell firing patterns, is the distribution of GABA_A_ subunits in the hippocampus. Receptor subunits are linked closely with specific frequencies due to their kinetics ^80^. Additionally different GABA_A_ subunits reside on different parts of the pyramidal cells in CA1, which means that it will affect CA3 and entorhinal inputs differently ^81^. Given that diazepam has higher affinity to α1-3, and α5 subunits, which are localized to the layers correlating with the entorhinal inputs, diazepam may be affecting the entorhinal inputs more than the CA3 inputs.

Diazepam is a positive allosteric modulator of GABA_A_, so the immediate effect is increased hyperpolarization of the cells it acts upon. Consistent with this, we found a reduction in spiking output on secondary neurons 10 ms after receiving inhibitory input (Fig. 2), suggesting that systemic diazepam is having a direct effect on the hippocampal circuit. However systemic diazepam could also affect regions beyond the hippocampus. Manipulations of the prelimbic cortex and medial septum both impact hippocampal function ^6,7,82–87^. Considering this, it is likely that systemic diazepam is producing both intra and extra hippocampal effects. Nevertheless, as noted above, diazepam is taken systematically as a drug by millions of people ^60^, so it is important to understand its effects on critical neural systems, like the hippocampus.

In humans, short term memory side effects of diazepam have been observed using paradigms such as the N-back, 3-word recall, or stop and go ^61–63^; however, these tasks are all measuring short-term and working memory, not long-term consolidation effects. SWRs are primarily thought to be involved in systems consolidation ^24,27,40,78,88–91^), in which information within the hippocampus and cortex change to better reflect the schema of the world^40,73,92^. Considering the massive disruption seen to essential hippocampal processes such as SWRs after systemic diazepam, further investigation should be made into long term memory and consolidation side effects, such as the development of environmental schema ^39^, task expertise ^93^, and as seen in _PTSD_ ^94–96^.

Given the effects of systemic diazepam on hippocampal function and representations, it is likely that diazepam has major disruptive effects to critical memory function, particularly effects on long-term consolidation. We suggest that further human studies are warranted, examining long-term consolidation effects rather than short-term working memory effects.

## Supplemental Figures and Tables

**Figure S1.**
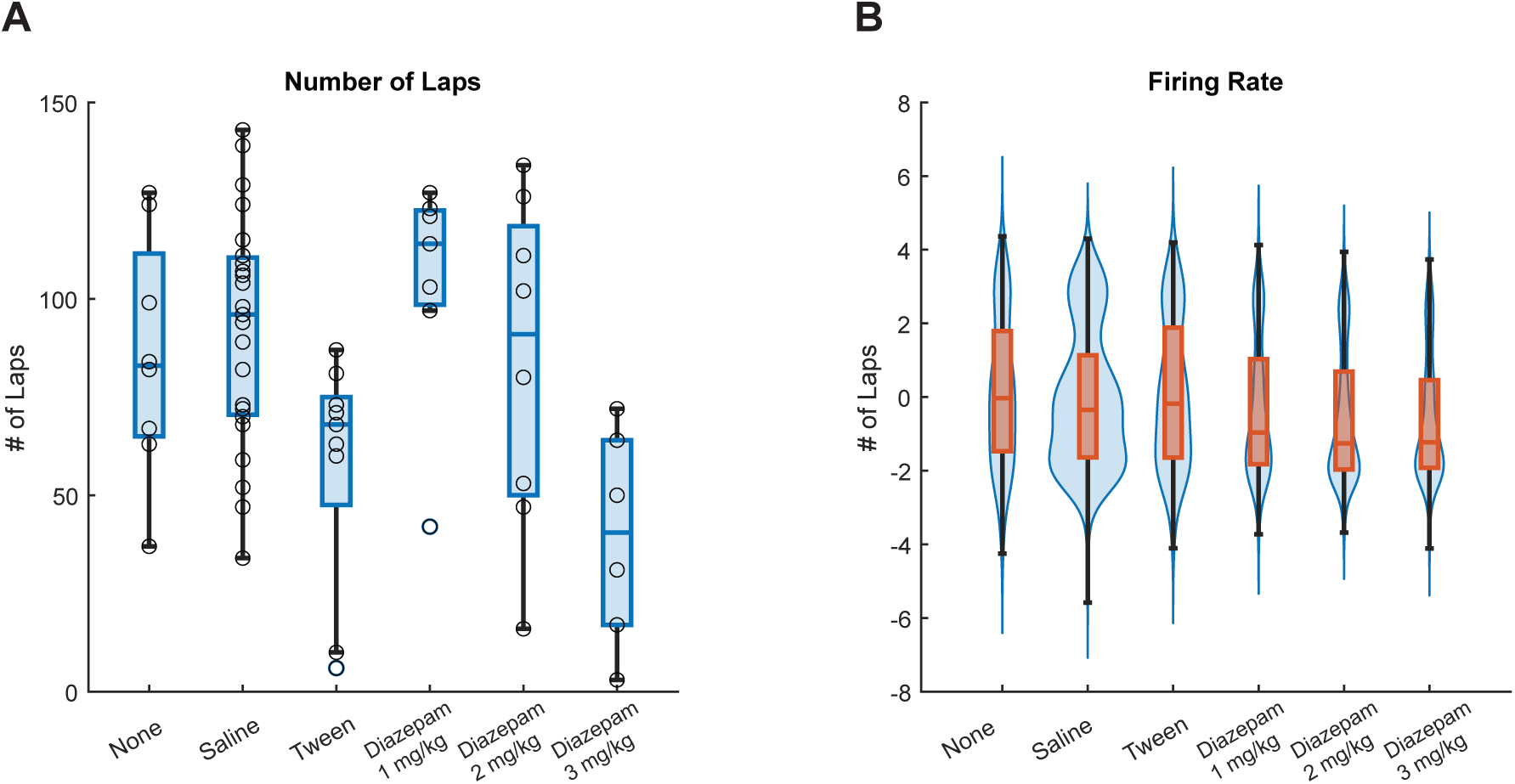
(A) Number of laps rats ran on average for no drug, saline, tween, and diazepam at 1, 2, and 3 mg/kg conditions. (B) Firing rate distribution of cells for no drug, saline, tween, and diazepam at 1, 2, and 3 mg/kg conditions.

**Figure S2.**
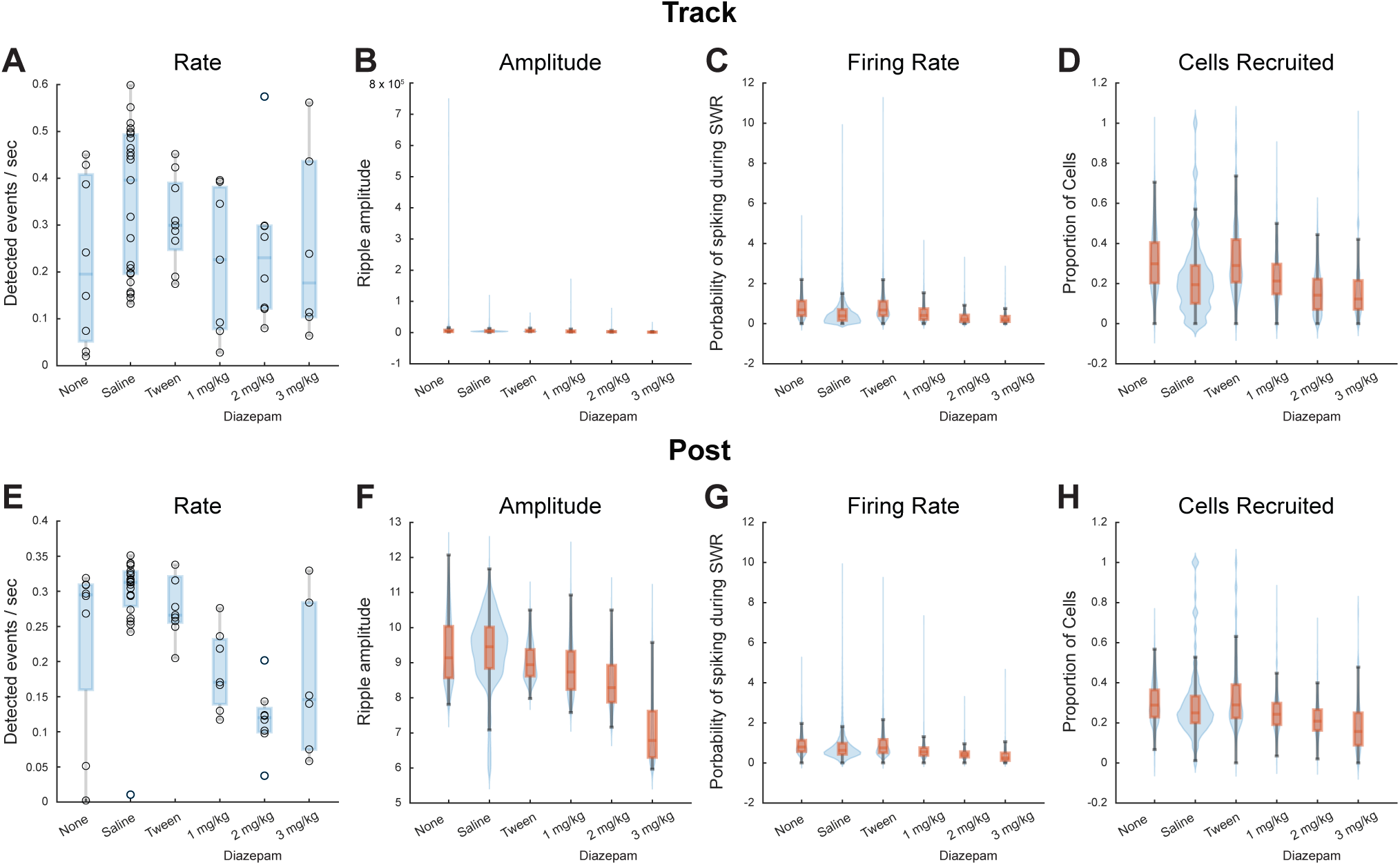
Sharp wave ripple (SWR) analyses for no drug, saline, tween, and diazepam at 1, 2, and 3 mg/kg conditions. (A,E) SWR rate. (B,F) SWR amplitude. (C,G) Firing rate during SWRs. (D,H) Proportion of cells firing within a given SWR. (A-D) On the track; (E-H) During post-run recording.

**Figure S3.**
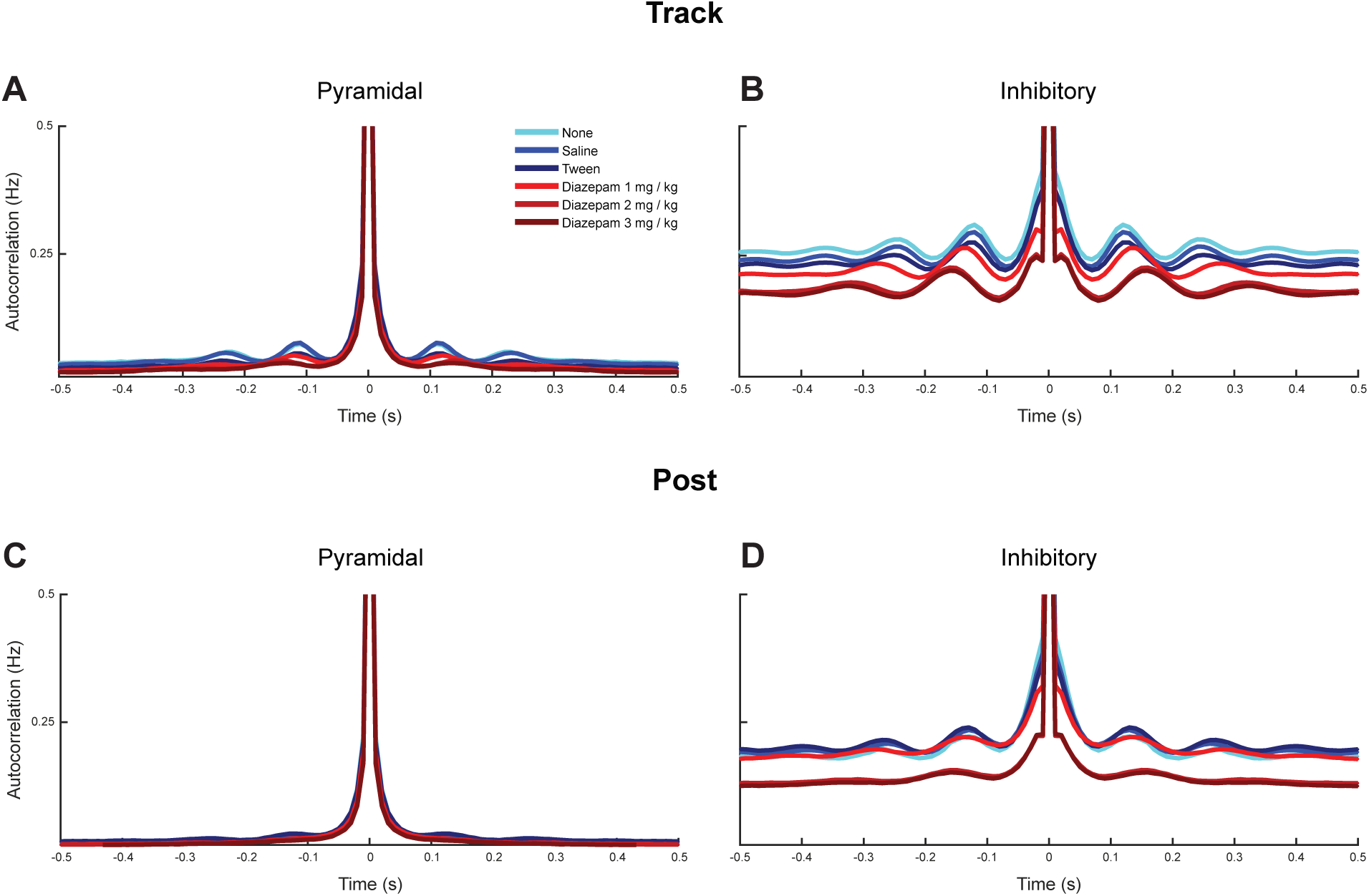
Spike autocorrelation analyses for no drug, saline, tween, and diazepam at 1, 2, and 3 mg/kg conditions. (A) Pyramidal neurons on the track. (B) Interneurons on the track. (C) Pyramidal neurons during post-run rest. (D) Inhibitory neurons during post-run rest.

**Figure S4.**
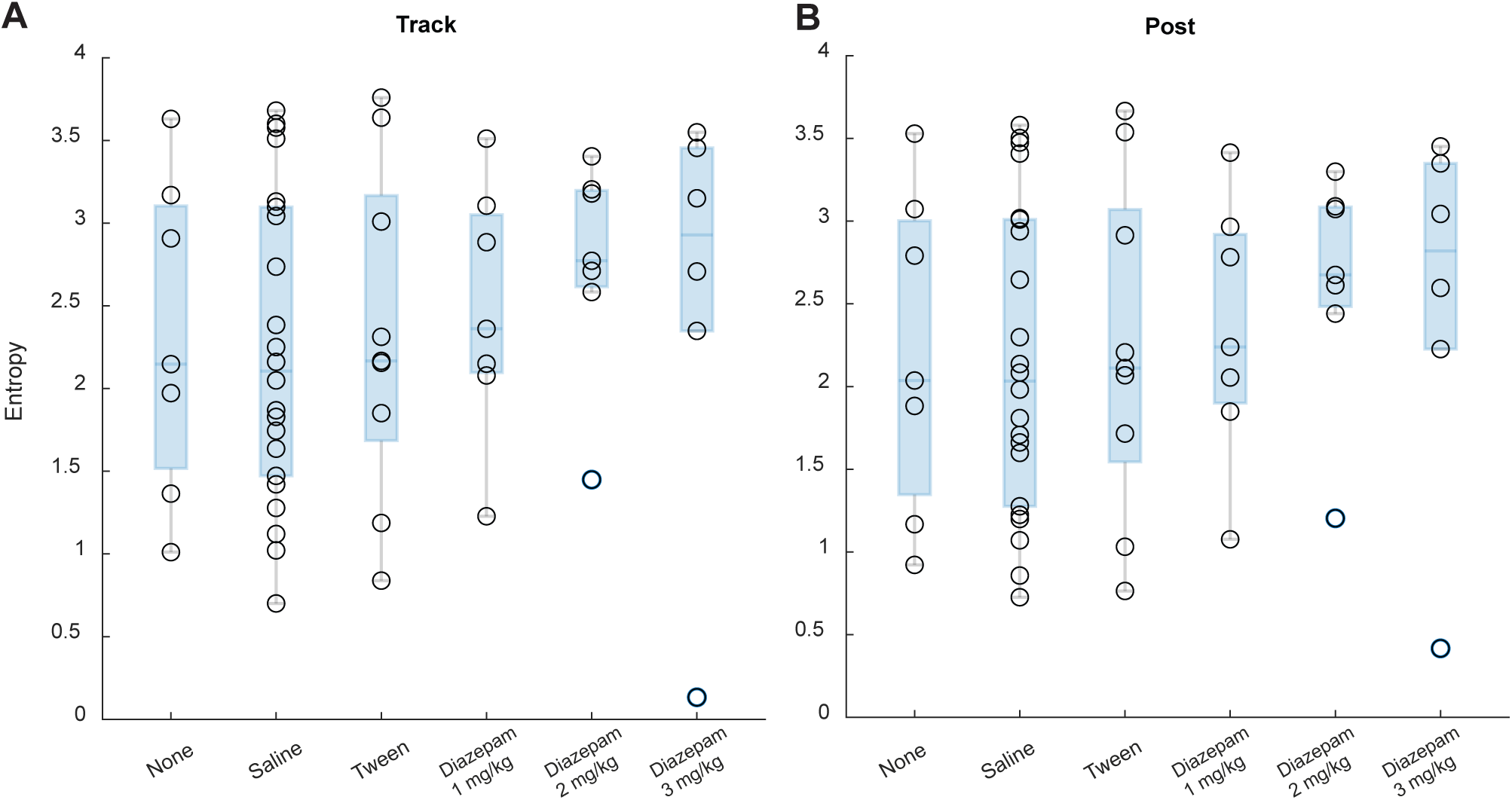
Average entropy of all decoding samples on the track (A) and during the post-run recording (B). No drug, saline, tween, and diazepam at 1, 2, and 3 mg/kg conditions shown. None of the differences are statistically significant.

**Table S1.**
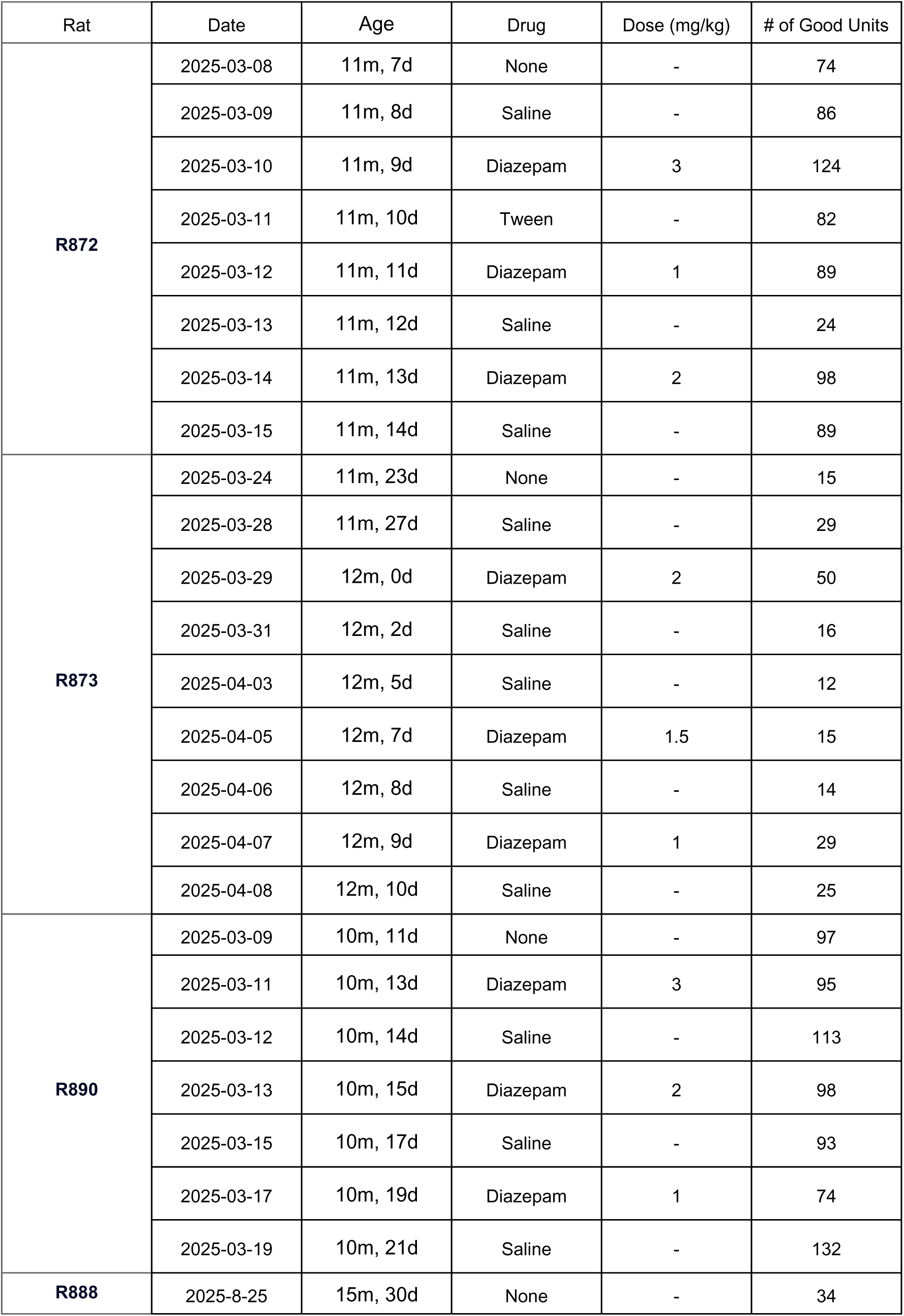
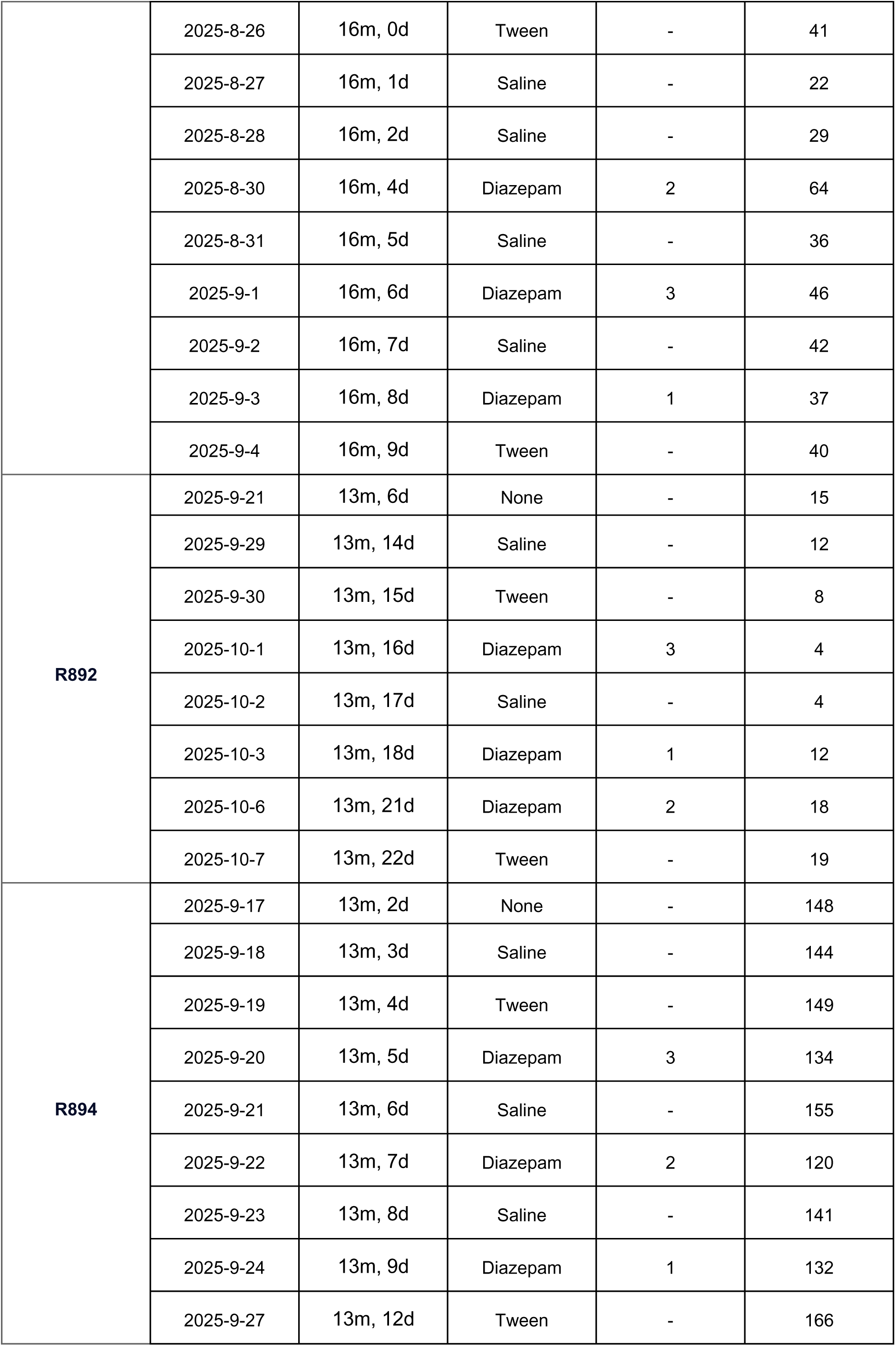
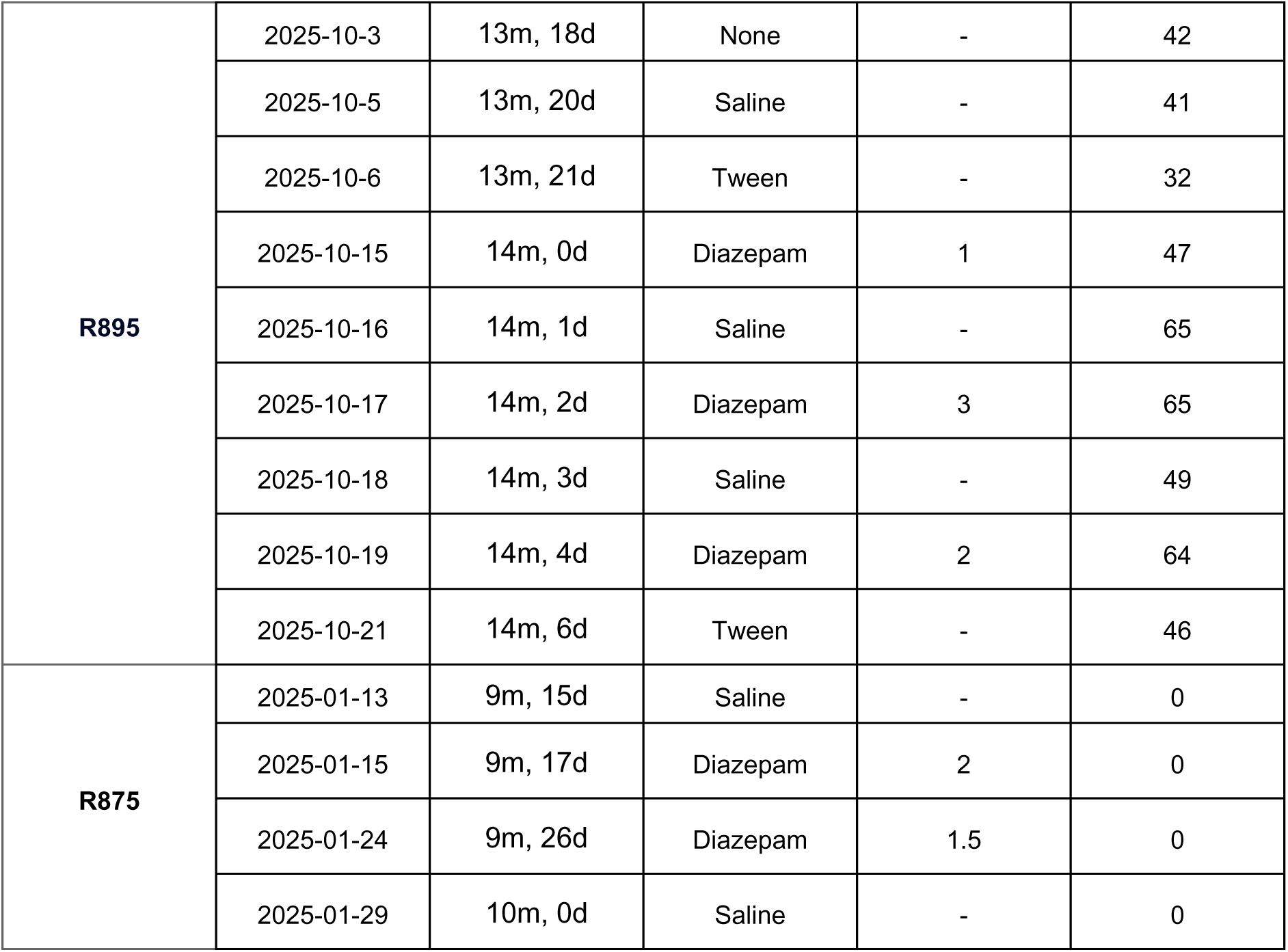
Rat ID (Column 1), date (Column 2), age of the animal (Column 3), drug condition (Column 4), dose in mg/kg (Column 5), and number of good units identified using Kilosort 2 and adjusted manually with Phy (Column 6).

| Rat | Date | Age | Drug | Dose (mg/kg) | # of Good Units |
| --- | --- | --- | --- | --- | --- |
| **R872** | 2025-03-08 | 11m, 7d | None | - | 74 |
|  | 2025-03-09 | 11m, 8d | Saline | - | 86 |
|  | 2025-03-10 | 11m, 9d | Diazepam | 3 | 124 |
|  | 2025-03-11 | 11m, 10d | Tween | - | 82 |
|  | 2025-03-12 | 11m, 11d | Diazepam | 1 | 89 |
|  | 2025-03-13 | 11m, 12d | Saline | - | 24 |
|  | 2025-03-14 | 11m, 13d | Diazepam | 2 | 98 |
|  | 2025-03-15 | 11m, 14d | Saline | - | 89 |
| **R873** | 2025-03-24 | 11m, 23d | None | - | 15 |
|  | 2025-03-28 | 11m, 27d | Saline | - | 29 |
|  | 2025-03-29 | 12m, 0d | Diazepam | 2 | 50 |
|  | 2025-03-31 | 12m, 2d | Saline | - | 16 |
|  | 2025-04-03 | 12m, 5d | Saline | - | 12 |
|  | 2025-04-05 | 12m, 7d | Diazepam | 1.5 | 15 |
|  | 2025-04-06 | 12m, 8d | Saline | - | 14 |
|  | 2025-04-07 | 12m, 9d | Diazepam | 1 | 29 |
|  | 2025-04-08 | 12m, 10d | Saline | - | 25 |
| **R890** | 2025-03-09 | 10m, 11d | None | - | 97 |
|  | 2025-03-11 | 10m, 13d | Diazepam | 3 | 95 |
|  | 2025-03-12 | 10m, 14d | Saline | - | 113 |
|  | 2025-03-13 | 10m, 15d | Diazepam | 2 | 98 |
|  | 2025-03-15 | 10m, 17d | Saline | - | 93 |
|  | 2025-03-17 | 10m, 19d | Diazepam | 1 | 74 |
|  | 2025-03-19 | 10m, 21d | Saline | - | 132 |
| **R888** | 2025-8-25 | 15m, 30d | None | - | 34 |
|  | 2025-8-26 | 16m, 0d | Tween | - | 41 |
|  | 2025-8-27 | 16m, 1d | Saline | - | 22 |
|  | 2025-8-28 | 16m, 2d | Saline | - | 29 |
|  | 2025-8-30 | 16m, 4d | Diazepam | 2 | 64 |
|  | 2025-8-31 | 16m, 5d | Saline | - | 36 |
|  | 2025-9-1 | 16m, 6d | Diazepam | 3 | 46 |
|  | 2025-9-2 | 16m, 7d | Saline | - | 42 |
|  | 2025-9-3 | 16m, 8d | Diazepam | 1 | 37 |
|  | 2025-9-4 | 16m, 9d | Tween | - | 40 |
| **R892** | 2025-9-21 | 13m, 6d | None | - | 15 |
|  | 2025-9-29 | 13m, 14d | Saline | - | 12 |
|  | 2025-9-30 | 13m, 15d | Tween | - | 8 |
|  | 2025-10-1 | 13m, 16d | Diazepam | 3 | 4 |
|  | 2025-10-2 | 13m, 17d | Saline | - | 4 |
|  | 2025-10-3 | 13m, 18d | Diazepam | 1 | 12 |
|  | 2025-10-6 | 13m, 21d | Diazepam | 2 | 18 |
|  | 2025-10-7 | 13m, 22d | Tween | - | 19 |
| **R894** | 2025-9-17 | 13m, 2d | None | - | 148 |
|  | 2025-9-18 | 13m, 3d | Saline | - | 144 |
|  | 2025-9-19 | 13m, 4d | Tween | - | 149 |
|  | 2025-9-20 | 13m, 5d | Diazepam | 3 | 134 |
|  | 2025-9-21 | 13m, 6d | Saline | - | 155 |
|  | 2025-9-22 | 13m, 7d | Diazepam | 2 | 120 |
|  | 2025-9-23 | 13m, 8d | Saline | - | 141 |
|  | 2025-9-24 | 13m, 9d | Diazepam | 1 | 132 |
|  | 2025-9-27 | 13m, 12d | Tween | - | 166 |
| **R895** | 2025-10-3 | 13m, 18d | None | - | 42 |
|  | 2025-10-5 | 13m, 20d | Saline | - | 41 |
|  | 2025-10-6 | 13m, 21d | Tween | - | 32 |
|  | 2025-10-15 | 14m, 0d | Diazepam | 1 | 47 |
|  | 2025-10-16 | 14m, 1d | Saline | - | 65 |
|  | 2025-10-17 | 14m, 2d | Diazepam | 3 | 65 |
|  | 2025-10-18 | 14m, 3d | Saline | - | 49 |
|  | 2025-10-19 | 14m, 4d | Diazepam | 2 | 64 |
|  | 2025-10-21 | 14m, 6d | Tween | - | 46 |
| **R875** | 2025-01-13 | 9m, 15d | Saline | - | 0 |
|  | 2025-01-15 | 9m, 17d | Diazepam | 2 | 0 |
|  | 2025-01-24 | 9m, 26d | Diazepam | 1.5 | 0 |
|  | 2025-01-29 | 10m, 0d | Saline | - | 0 |

## Materials and Methods

### Experimental Design and Statistical Analyses

#### Subjects

Seven female and one male Fisher Brown Norway rat aged 9-11 months were the subject of the experiment. Rats were kept in a 14:10 h light/dark cycle in the vivarium. During the task, rats were food restricted and received 45 mg food pellets such that they maintained above 80% of their free-feeding weight. The reanalyzed cohort providing the data used for Fig. 5 consisted of six male and five female Brown Norway rats aged 8-13 months run on an approach-avoidance motivational conflict task ^44^. Rats were maintained in the same conditions as described above. One day from one rat (ID R554), was not included in analysis due to poor tracking. All experimental procedures were approved by the University of Minnesota Institutional Animal Care and Use Committee (IACUC). All aspects of both experiments were performed in alignment with NIH guidelines.

#### Surgery

Rats were initially anesthetized with 5% isoflurane in 1 L of oxygen. After initial anesthetic, rats were secured to the stereotaxis and maintained at 1-2% isoflurane in 1 L of oxygen. Respirations, temperature, and reaction were monitored throughout the procedure following IACUC guidelines. Silicon probes were chronically implanted through a burr hole created in skull bone. The dura was removed before probe insertion. Probes were implanted at −3.8 mm anterior-posterior and (+/−)2.5 mm medial-lateral. Craniotomy was sealed with bone wax and a 3D printed skull ring and housing was cemented to the skull via Metabond. Rats received 0.8 mL of Children’s Tylenol 5-10 minutes after being taken off stereotaxic. Baytril and carprofen were administered according to rat weight for 3 days after surgery. Betadine was used to clean around the implant for 7 days after surgery. On post-op day 3, rats were placed on food restriction and began training the day after.

The reanalyzed cohort providing data used for Figure 5 was implanted with 64 channel H3 probes (Cambridge Neurotech) except for one rat who was implanted with a 64 channel E2 probe (Cambridge Neurotech). One rat (ID R551) was excluded from the data set due to incorrect targeting.

#### Pharmacology

The primary cohort of rats were administered a range of diazepam doses including 1 mg/kg, 1.5 mg/kg, 2 mg/kg, and 3 mg/kg. As only one rat was administered the 1.5 mg/kg dose, it was not analyzed here. All rats received 1 mg/kg, 2 mg/kg and 3 mg/kg doses. A stock solution was created by dissolving diazepam into Tween-20. The stock solution was diluted with 0.9% saline to reach target concentration. For control manipulations, we included both Tween-20 vehicle solution and saline only injection equivalent to the volume of diazepam administered. Diazepam and Tween-20 were sourced from Sigma Adlrich. The drug and controls were administered via intraperitoneal injection (IP) 10 minutes prior to the task.

#### Experimental Design

The linear track was 32 cm wide and 112 cm long, constructed of lego bricks of random color with 3D printed plastic feeders at each end of the track. Rats were first trained to alternate for three 45mg full-nutrition food pellets (5TUL, TestDiet) at each end, which was then reduced to two 45mg pellets once they were running laps reliably. During training, rats ran on the linear track for 30 minutes. The behavior analyzed is after training, and consisted of 30 minute sessions on the linear track, receiving 2 pellets at each end, followed by a 20 minute post-run recording. The 20 minute post-run recording was only included for recording, not training days. After learning the task, control and diazepam days were alternated, but were always separated by at least one day in which the rat ran the track with no injection or was fed its daily food from a ramekin in a separate cage in between. Details on the behavioral experimental design for the second (reanalyzed) cohort (Figure 5) are available from ^44^.

#### Code / Software

Behavior was identified using tracking from an LED attached to the rat’s headstage. For each session, the “best” LFP was identified from one channel in layer by visually identifying the least noisy frequency-frequency correlation plots ^45^. Cells were identified using Kilosort v2 and then manually adjusted using Phy. Cells with a coefficient of variation of the interspike intervals ^2^ (CV2) > 1 and firing rate < 3 Hz were identified as pyramidal neurons, while cells with CV2 < 1 or firing rate > 3 Hz were identified as interneurons.

SWRs were detected by first filtering and Hilbert transforming the LFP in the ripple power range (140-200 Hz), and then applying a threshold of z > 2.5. Amplitude was determined at the peak of the SWR. To examine cellular activity, firing rate was calculated as the average of each cell’s spike count by duration of the SWR. Calculated firing rate was averaged across all detected SWRs.

Theta was defined as 6-10 Hz, lo-gamma 30-50 Hz, and hi-gamma 80-120 Hz. Phase and amplitude were identified through filtering and Hilbert transforms.

Phase coupling was calculated per the pairwise phase consistency (PPC) which compares the angles between all spike pairs to matched shuffles ^46,47^

Decoding was performed using a one-step Bayesian decoder from spatial tuning curves of the animal’s position on the linear track ^28,29^ using 64 linear bins along the 13.75 x 24 inch track with a time bin of 250 ms.. Decoded posterior probabilities were used to calculate the entropy of the decoding.

Rat ages, drug dose sequence, and recording cell yields are reported in Supplemental Table 1.

## Conflict of Interest

The authors declare no competing financial interests.

## Acknowledgments

Funded by R01-MH080318, R01-MH112688, the University of Minnesota, including the Undergraduate Research Opportunities Program (UROP, University of Minnesota). Special thanks to the Faculty for Undergraduate Neuroscience (FUN) and International Brain Research Organization (IBRO) for funding the first author’s travel to present this work at the Society for Neuroscience (SFN). Thank you to Chris Boldt and Celia Gagliardi for help with this project.

## Notes

### Competing Interest Statement

The authors have declared no competing interest.

